# Network Modeling and Inference of Peroxisome Proliferator-Activated Receptor Pathway in High fat diet-linked Obesity

**DOI:** 10.1101/2020.09.15.298356

**Authors:** Haswanth Vundavilli, Lokesh P. Tripathi, Aniruddha Datta, Kenji Mizuguchi

## Abstract

Systems biology aims to understand how holistic systems theory can be used to explain the observable living system characteristics, and mathematical modeling tools have been successful in understanding the intricate relationships underlying cellular functions. Lately, researchers have been interested in understanding molecular mechanisms underlying obesity, which is a major health concern worldwide and has been linked to several diseases. Various mechanisms such as peroxisome proliferator-activated receptors (PPARs) are known to modulate obesity-induced inflammation and its consequences. In this study, we have modeled the PPAR pathway using a Bayesian model and inferred the sub-pathways that are potentially responsible for the activation of the output processes that are associated with high fat diet (HFD)-induced obesity. We examined a previously published dataset from a study that compared gene expression profiles of 40 mice maintained on HFD against 40 mice fed with chow diet (CD). Our simulations have highlighted that GPCR and FATCD36 sub-pathways were aberrantly active in HFD mice and are therefore favorable targets for anti-obesity strategies. We further cross-validated our observations with experimental results from the literature. We believe that mathematical models such as those presented in the present study can help in inferring other pathways and deducing significant biological relationships.

## 1 Introduction

The remarkable increase in omics data such as those in genomics, proteomics, and metabolomics, has revolutionized biological research and led to an emergence of new disciplines and methodologies that are able to study complex biological problems in an unprecedented manner [1]. These research developments have been primarily possible because of the proliferation of technologies that can reliably measure the concentrations of various cellular components and also their interactions [2]. Specifically, advancements in genomics have launched molecular biology into the realms of systems biology, which integrates computational and experimental research to define relationships between genes, pathways and phenotypes [3]. Lately, researchers have developed and applied mathematical models to study various biological systems such as cancer [4]. Such models have been widely employed to examine the perturbations of bio-molecular relationships in disease biology and they have shown significant successes in contributing to successful therapeutic outcomes [5].

Obesity is linked with several chronic conditions such as cardiovascular diseases and other major health concerns [6]. Despite the recent strides in research exploring the causality of these links, much of the molecular mechanisms underlying obesity remain only vaguely understood. However, recent advances in omics have made it possible to obtain large-scale datasets which when analyzed using mathematical techniques such as network inference [7], can pinpoint specific cellular factors, sub-pathways and subnetworks the perturbation of which can be associated with specific diseases.

One approach to studying the fundamental connections in the context of obesity is to critically examine the perturbations in the underlying metabolic pathways that maybe responsible for obesity and the related phenotypes. In this study, we have examined the data from a previously published study that investigated the effects of high-fat diet (HFD) on mitochondrial functioning in liver metabolism. The dataset comprised of transcriptomics profiles of 80 mice cohorts that were maintained on different dietary regimes: chow diet (CD) (6% kcal of fat) and HFD (60% kcal of fat) [8]. In the study, the pathways associated with fatty acid metabolism and storage, in particular, the peroxisome proliferator-activated receptor (PPAR) pathway, emerged as highly enriched and the most variable across CD and HFD cohorts. PPARs are nuclear receptor proteins and oversee key roles in cellular development, differentiation, and metabolism [9]. Therefore, we focused our efforts on investigating any potential HFD-induced aberrations in the PPAR pathway and their probable impact on the resulting phenotypes.

The paper is organized as follows. In Section 2, we first provide a review of Bayesian networks (BN) and derive the posterior probabilities. In Section 3, we reconstructed the PPAR pathway using KEGG and published literature. In Section 4, we then present our simulations and theoretical findings. Finally, in Section 5, we discuss the significance of our model in the context of HFD-induced obesity along with concluding remarks.

## 2 Methodology

Regulatory networks encompass multiple interactions among various biomolecules that underlie the functioning of key cellular processes [10]. Generally, these processes are well-regulated and tightly-controlled; network perturbations and loss of specific interactions within these processes are often linked with specific diseases. Over the years, computational tools and mathematical models have been widely and successfully used to model and interpret such complex and multi-layered networks from multi-dimensional omics data [11]. One such widely used tool is Bayesian Network (BN) modeling. We will now briefly review BNs and then outline our methodology in detail.

### Bayesian Networks

The interactions among the genes in a gene regulatory network are known to be relatively sparse. In other words, each gene within a network interacts with a relatively smaller number of genes, as opposed to the total number of genes within that network. This sparseness makes BN modeling an attractive option for simulating gene regulatory networks [12]. A bayesian network is represented by 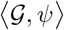, where 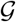 is a Directed Acyclic Graph (DAG) and *ψ* is the Conditional Probability Distribution (CPD) of each node. Each node in 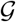 represents a random variable and the edges represent dependencies between the nodes.

Using the local Markov independence assumption, any Joint Probability Distribution (JPD) that satisfies the Markov condition can be described as a product of CPDs [3]. That is,

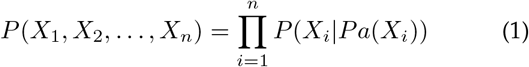

where *X_i_* is a random variable and *Pa*(*X_i_*) is the set of parents of *X_i_*. In the next section, we describe the construction of the graph structure 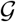 for the PPAR signaling pathway based on KEGG and published literature; this graph structure forms the basis for the factorization of the JPD of the model. We will then used gene expression data to update the model parameters *ψ*.

### Discretizing the gene expression data

For each gene, we discretized the expression values into a binary framework. Quantizing the gene expression values into a binary format helps to reduce computational complexity and in enhancing the robustness against noise [13]. Using the maximum likelihood estimator for the mean *μ* as the threshold, we discretized the expression data for each gene, where values above a threshold were assigned a value of ‘1’, and those below the threshold were assigned a value of ‘0’. This threshold is justified by the central limit theorem and the law of large numbers.

### Prior and Posterior distributions

Next, we used bayesian modeling to estimate the network parameters. Let *ψ_X_* be the probability that a variable *X* takes the value ‘1’. The prior of an uncertain quantity, such as *ψ_X_*, is a probability distribution of the quantity before the evidence is considered. For each variable, we assumed that *ψ_X_* was Beta distributed. That is,

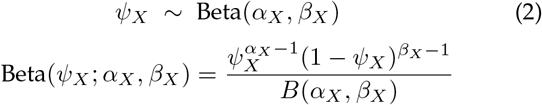

where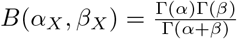 and Γ is the standard Gamma function.

The data likelihood for *n* observations is defined as:

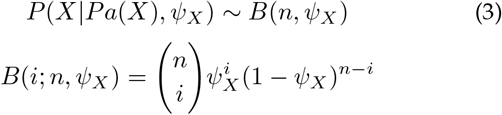

where *i* is the number of successes (‘1’s).

Since the binomial likelihood is a conjugate to the beta distribution, the posterior distributions of the variables were defined as:

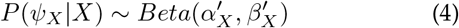

where 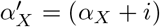 and 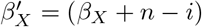.

Using the above equation, the expected value can then be defined as:

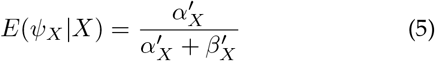

Similarly, if we have two nodes *X* and *Y*, connected such that *Y* is the parent of *X*, the conditional posterior probability is defined as:

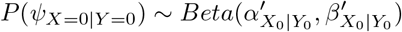

and the expected value of *X*_0_|*Y*_0_ is defined as:

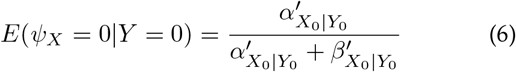

where 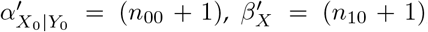, *n*_00_ is the number of observations where both *X* and *Y* are ‘0’s, and *n*_00_ is the number of observations where *X* is ‘1’ and *Y* is ‘0’ simultaneously.

### Computing probabilities

Using the above equations, we may now compute the probability that a gene is over-expressed as the ratio of over-expressed samples (number of ‘1s) to the total number of samples in that dataset. That is,

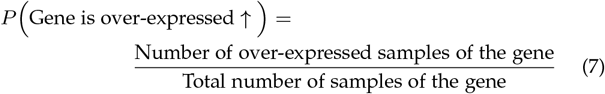

Similarly, the probability that a gene is under-expressed may be computed as the ratio of under-expressed samples (number of ‘0s) to the total number of samples in the dataset. That is,

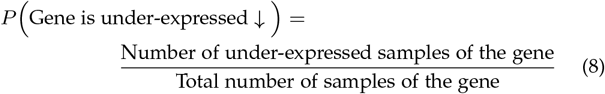

We may also compute the conditional probability that a gene *G*_1_ is over-expressed given a gene *G*_2_ upstream is over-expressed simultaneously as:

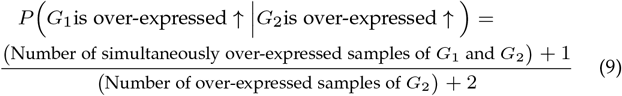

Similarly, the conditional probability that a gene *G*_1_ is under-expressed given a gene *G*_2_ upstream is underexpressed simultaneously can be computed as:

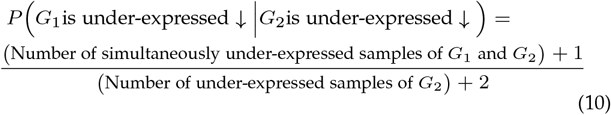

We will now briefly discuss the PPAR pathway and apply the above methodology to infer the aberrant sub-pathways.

## 3 PPAR Pathway

Peroxisome proliferator-activated receptors (PPARs) are ligand-activated transcription factors and are a part of the nuclear hormone receptor family. They occur as three isoforms: PPAR*α*, PPAR*β/δ*, and PPAR*γ* [14]. Each subtype functions in a different manner probably because of their distinctive tissue distributions and biochemical properties.

PPARs are stimulated by fatty acids or their derivatives and play decisive roles in multiple biological processes, including lipid metabolism, glucose metabolism, and overall energy homeostasis. PPARs form heterodimers with retinoid X receptor (RXR) and the resultant transcription factors regulate various cellular functions [15]. For example, PPAR*α*-RXR dimers stimulate genes that control lipogenesis and cholesterol metabolism, while PPAR*β/δ*-RXR dimers supervise ubiquitination and cell survival [16].

PPAR pathway is a complex signaling network and its dysregulation is linked with multiple diseases including liver cancer and fatty liver disease [17]. Unsurprisingly, PPAR pathway has been associated with obesity-induced inflammation making it an attractive therapeutic target for mitigating obesity and the associated health concerns. Hence, in this paper, we have investigated the highly aberrant sub-pathways within the PPAR pathway using mathematical modeling and we have inferred the most promising targets for therapeutic intervention to mitigate obesity.

Figure 1 illustrates the PPAR pathway reconstructed from literature. The upstream nodes represent the genes and the downstream nodes represent the output processes.

**Fig. 1:**
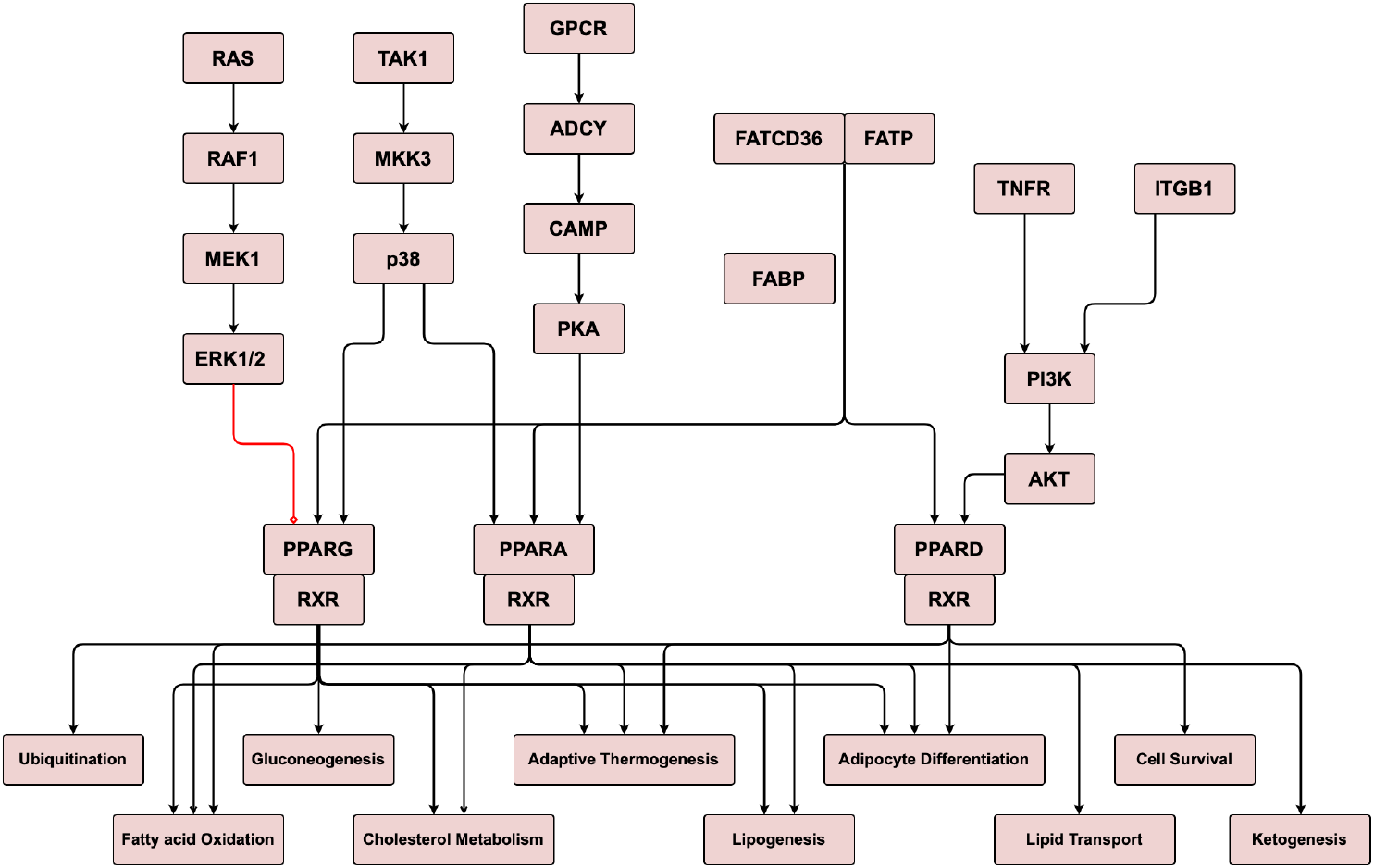
Peroxisome proliferator-activated receptors pathway (PPAR). Upstream nodes represent the genes and the downstream nodes represent the output processes. Black arrows indicate activation and red arrows indicate inhibition.

Next, we applied our approach to a dataset of gene expression profiles from 40 mice fed with HFD and 40 mice fed with CD. Fundamentally, we aimed to identify differentially expressed genes (DEGs) and the affected sub-pathways in the PPAR pathway that were significantly influenced by HFD.

## 4 Results

For each gene in the PPAR pathway, we first discretized the gene expression values. Then, using equation (7), we computed the probabilities of over-expression separately for HFD and CD samples. We then calculated the ratio of the over-expressed probabilities for each gene and plotted them as a bar chart in figure 2. In the figure, red bars imply a ratio greater than 1 and blue bars imply a ratio less than 1. The higher number of red bars clearly indicated that a majority of genes in the PPAR pathway were significantly over-expressed in HFD as compared to CD.

**Fig. 2:**
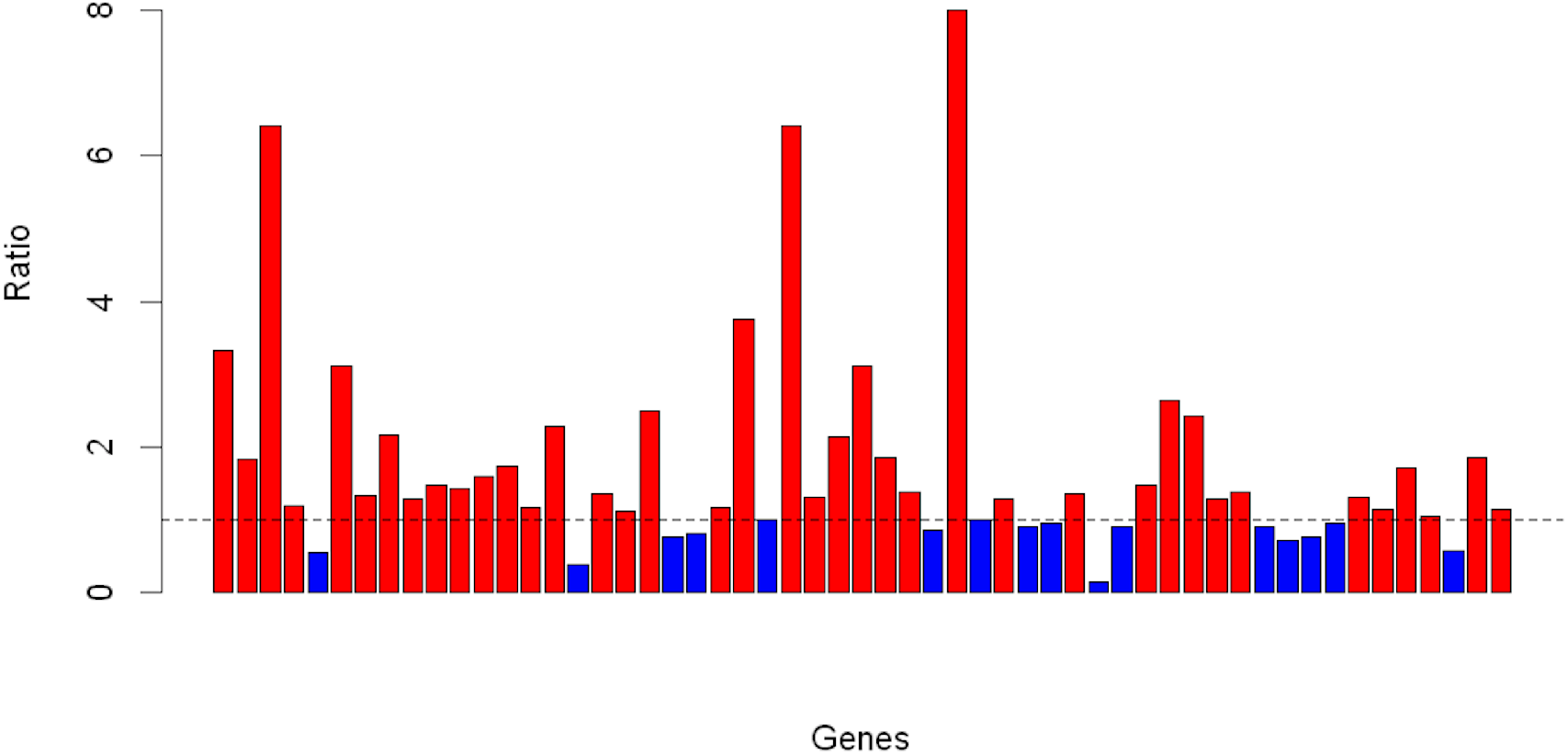
Ratio of over-expressed probabilities of genes in HFD samples to CD samples in the PPAR pathway (Red = Ratio > 1, Blue = Ratio < 1).

Next, we investigated if the same assertion held true for the output processes, that is, whether the output processes were also significantly activated in HFD as compared to CD. We modeled the output processes using the same approach and plotted the ratio of the probabilities of output processes in HFD compared to CD as summarized in figure 3. As before, the red bars imply a ratio greater than 1 and blue bars imply a ratio less than 1; evidently, the output processes were also significantly activated in HFD as compared to CD.

**Fig. 3:**
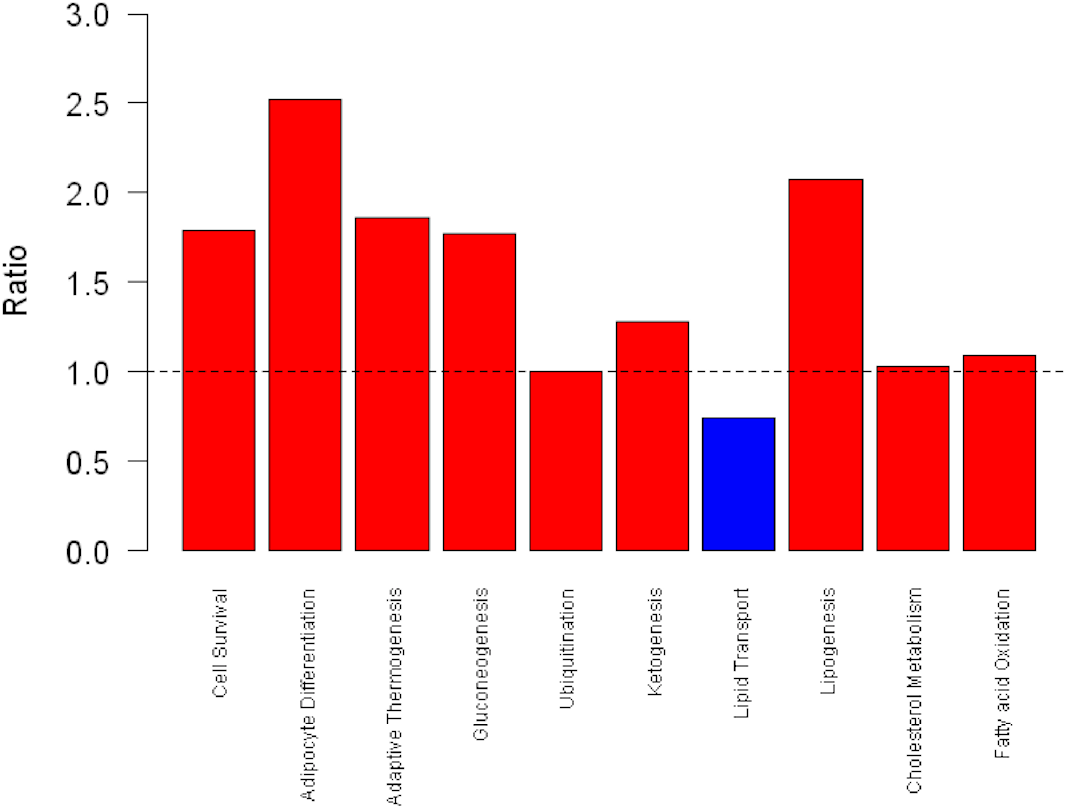
Ratio of over-expressed probabilities of output processes in HFD samples to CD samples in the PPAR pathway (Red = Ratio > 1, Blue = Ratio < 1).

To obtain an overview of the genes and sub-pathways responsible for the over-expression of genes and output processes in HFD, we overlaid these probabilities onto the PPAR pathway. A probability of less than 0.4 was highlighted in blue, a probability less than 0.6 but greater than 0.4 was highlighted in light-red, and a probability greater than 0.6 was highlighted in dark-red. In figure 4, we overlaid the probabilities and the corresponding colors onto the PPAR pathway for HFD and CD. As expected, the genes and processes in HFD were mostly dark-red (over-expressed) while CD were mostly blue (under-expressed). It also became evident that lipogenesis and adipocyte differentiation were the most perturbed output processes that were strongly activated in HFD compared with CD (Figures 3, 4). These two processes have also been shown to be abnormally active in previous studies thereby, corroborating our observations [18], [19].

**Fig. 4:**
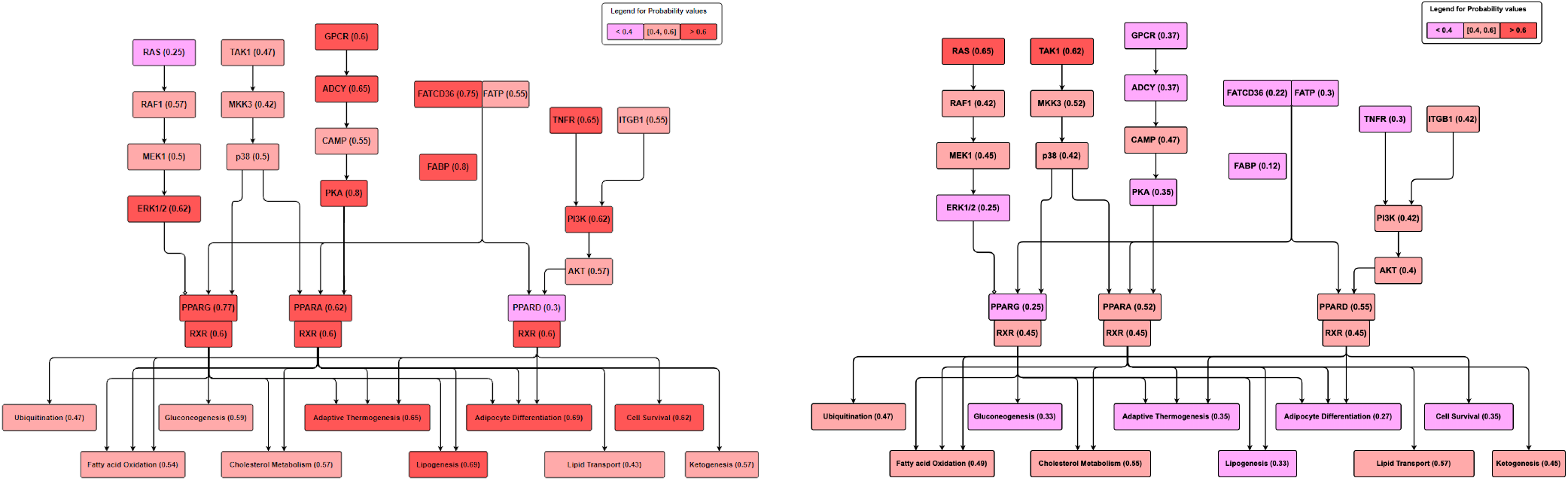
Probability that a gene/process is over-expressed/activated in high fat diet (left) and chow diet (right). The genes and processes in HFD samples are mostly dark-red (over-expressed) while in CD samples are mostly blue (under-expressed).

To understand the sub-pathways that were likely responsible for the hyper-activation of lipogenesis and adipocyte differentiation, we computed the conditional probabilities of the gene-gene interactions using equations (9) and (10). Then, we scanned for sub-pathways that included aberrantly expressed genes as well as high gene-gene conditional probabilities. This cross-validation was especially relevant since we are interested in mapping the interaction flow that was potentially leading up to the aberrations in these output processes. Hence, we overlaid the conditional probabilities for both HFD and CD samples on the PPAR pathway and highlighted the gene-gene interaction flows for lipogenesis and adipocyte differentiation using red arrows (Fig. 5). From the figure, GPCR and FATCD36 subpathways were strongly activated in HFD as compared to CD, and therefore, it is likely that these sub-pathways have the maximal impact on the two output processes.

**Fig. 5:**
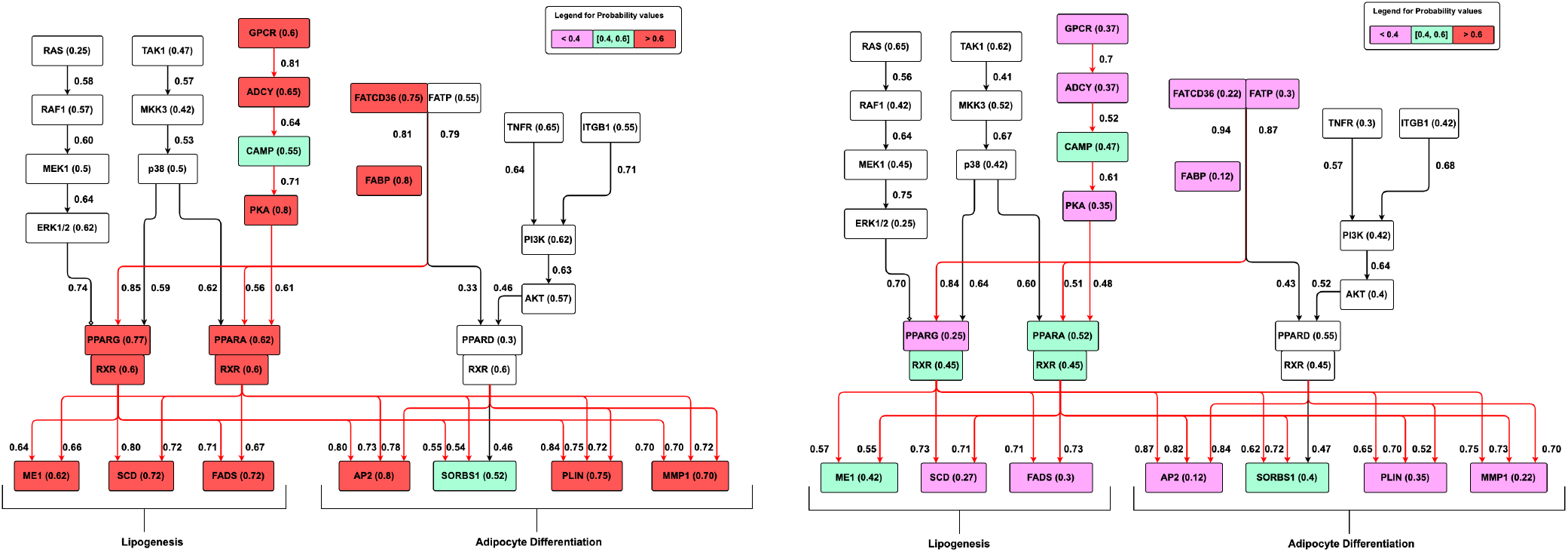
The sub-pathways that were perturbed in HFD (left) and CD (right). The sub-pathways are strongly active in HFD as compared to CD.

After highlighting the most relevant sub-pathways, we next examined the seemingly most suitable targets that could be potentially manipulated to restore any aberrations in these processes to their ‘normal’ states. To pinpoint suitable target genes for such purposes, we computed the conditional probability that a gene was under-expressed given that a gene upstream was under-expressed simultaneously supposedly through intervention or therapy as in equation (10). The conditional probabilities were then assembled into a heatmap in figure 6. Genes highlighted in shades approaching blue were deemed to be better therapeutic targets as compared to those with shades approaching red. For each output process, we also averaged the values to identify the best overall target for that process. As is evident from the heatmap, PPARA, a regulator of lipid homeostasis that has been previously targeted for hyperlipidemia treatments [20], emerged as the best overall target for adipocyte differentiation. In a recent study, PPARA was also shown to be neuroprotective in retinopathy of type 1 diabetes, and therefore is a potential therapeutic target for diabetic retinopathy [21]. G-protein coupled receptors (GPCRs) are a family of proteins that are responsible for multiple cellular responses by activating internal signal transduction pathways; they are also known to be important drug targets in various diseases [22]. Specifically, the leucine-rich repeat-containing G protein-coupled receptor 6 (LRG6) is a GPCR gene that activates the gene adenylate cyclase (ADCY) and eventually PPARA. From the heatmap, we inferred that the GPCR subpathway was the best overall target for lipogenesis.

**Fig. 6:**
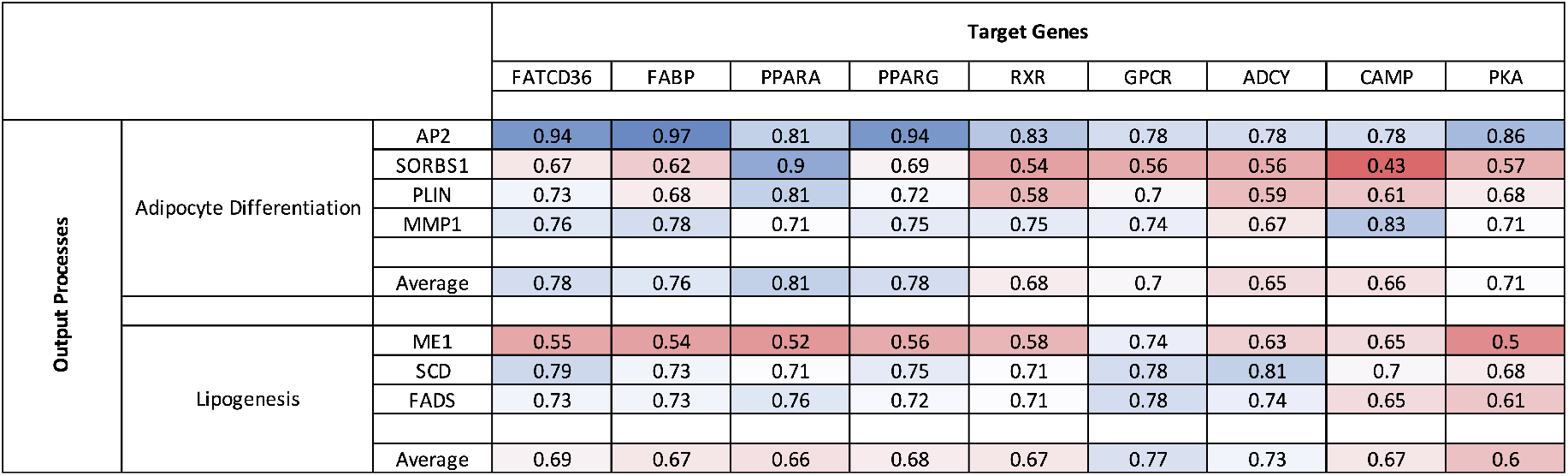
Heat map of the conditional probabilities of a gene being under-expressed given that a gene upstream is under-expressed simultaneously. A shade closer to blue is a better therapeutic target as compared to a shade closer to red.

## 5 Discussion

Advancements in computational techniques and systems biology have prompted many researchers to approach biological processes as control problems. The aim of these control problems is to design engineering methods to steer an abnormal biological network into an acceptable state by a suitable therapeutic intervention [23]. There are multiple advantages to this approach, since a realistic and sufficiently able model will be able to reliably predict and rank therapeutic outcomes without having to rely solely on costly, laborious and time consuming experiments. For example, a typical biological network with five genes would require about 31 different experiments to determine the best combination of target(s) for that network. Mathematical tools can help in reducing the search space by performing meaningful simulations.

The PPARs and the RXRs form heterodimers and collectively regulate gene expression and perform important roles in various cellular processes such as fatty acid storage, glucose homeostasis, and energy balance. Abnormalities or dysfunctions in any of these processes are associated with diseases such as obesity and diabetes [24]. Our observations have therefore, strongly supported the suitability of the PPAR pathway as a potential target for therapeutic intervention in obesity. Hence, obtaining deeper biological insights into the functioning of the PPAR pathway may help uncover key sub-pathways and potentially novel targets for newer and more effective therapeutic applications.

In this study, we have presented a bayesian framework to infer sub-pathways within the PPAR pathway that were likely influenced aberrantly under different dietary scenarios. Our simulations showed that GPCR and FATCD36 subpathways were aberrantly active in HFD as compared to CD. Our observations also agreed with those published elsewhere, thereby bolstering our model [25], [26]. We further examined the strongly activated output processes and prioritized the most likely therapeutic gene targets that may be potentially leveraged to restore these abnormal processes back to their normal states.

We conclude that our observations are in agreement with the previously published experimental results, thereby demonstrating the efficacy of our approach. We believe that our model can form the basis for developing new techniques and applying them to new complex biological networks.

## Acknowledgment

This work was supported in part by Grants-in-Aid for Scientific Research from the Japan Society for the Promotion of Science under Grant Number 17K07268 to K.M. This work was also supported in part by the US National Science Foundation under Grant ECCS-1609236 and in part by the TEES-AgriLife Center for Bioinformatics and Genomic Systems Engineering (CBGSE) startup funds to A.D. The statements made herein are solely the responsibility of the authors. Declarations of interest: none.

